# To improve the predictions of binding residues with DNA, RNA, carbohydrate, and peptide via multiple-task deep neural networks

**DOI:** 10.1101/2020.02.11.943571

**Authors:** Zhe Sun, Shuangjia Zheng, Huiying Zhao, Zhangming Niu, Yutong Lu, Yi Pan, Yuedong Yang

## Abstract

**Motivation:** The interactions of proteins with DNA, RNA, peptide, and carbohydrate play key roles in various biological processes. The studies of uncharacterized protein–molecules interactions could be aided by accurate predictions of residues that bind with partner molecules. However, the existing methods for predicting binding residues on proteins remain of relatively low accuracies due to the limited number of complex structures in databases. As different types of molecules partially share chemical mechanisms, the predictions for each molecular type should benefit from the binding information with other molecules types.

**Results:** In this study, we employed a multiple task deep learning strategy to develop a new sequence-based method for simultaneously predicting binding residues/sites with multiple important molecule types named MTDsite. By combining four training sets for DNA, RNA, peptide, and carbohydrate-binding proteins, our method yielded accurate and robust predictions with AUC values of 0.852, 0836, 0.758, and 0.776 on their respective independent test sets, which are 0.52 to 6.6% better than other state-of-the-art methods. More importantly, this study provides a new strategy to improve predictions by combining multiple similar tasks.

**Availability:** http://biomed.nscc-gz.cn/server/MTDsite/

## 1. Introduction

Predicting proteins interactions with other molecules is critical for understanding biological processes and discovering drugs. The most important molecules controlling biological activities include DNA, RNA, peptides and carbohydrates (CBH) (Hanson, et al., 2019). For example, interactions between protein and nucleic acids are central to many of the vital processes in molecular biology (Jia, et al., 2019) such as transcription, translation, post-transcriptional modification and regulation. Additionally, RNA-binding proteins can modulate or stabilize RNA structures to make RNA catalytically active (Pan and Shen, 2017). The carbohydrate-binding proteins act as important biomarkers for cell communication, cell adhesion, fertilization, development and differentiation (Lu and Pieters, 2019). Peptide interactions with protein domains occur in many cell processes particular signaling pathways (Petsalaki and Russell, 2008). In order to better understand the molecular mechanisms, we need to know the binding residues of these proteins interacting with their respective binding partner. Traditional experimental techniques such as X-ray and NMR experiments are robust but are expensive and time-consuming (London, et al., 2010) (Su, et al., 2018). With the exponentially increasing protein sequences, it is demanding to make predictions of binding residues from sequences.

Many bioinformatics methods combining physicochemical and evolutionary features (Wang L, 2010) have been developed in the past decades. For DNA-binding residue predictions, representative methods include TargetDNA (Jun Hu, 2017) and HMMBinder (Rianon Zaman, 2017) based on SVM, CNNsites (Wang, 2016) based on CNN network, DRNApred (Jing Yan, 2017) for accurately predicting and discriminating between DNA-and RNA-binding residues, and SPOT-DNA-Seq (Zhao, et al., 2014) based on alignments with known DNA-binding proteins. For RNA-binding residue predictions, there are RNAProSite (Meijian Sun, 2016) based on the random forest classifier and PredRBR (Yongjun Tang, 2017) based on the gradient tree boosting. For predictions of carbohydrate binding residues, the common idea is to find residues frequently observed on the sugar interface(Ghazaleh Taherzadeh, 2016) (Sujatha M S, 2004). For predictions of peptide-binding residues, several methods have been developed based on machine learning techniques, e.g. SPRINT(Zhou, 2016) and SPOT-Peptide(Zhou, 2019).

Although many successful methods have been proposed, most of them suffer from low accuracies due to small training sample sizes. The underlying reason is that the complex structures of proteins are difficult to obtain by experiments. To be worse, the predictions of binding residues with different types of molecules were traditionally treated as different problems (Miao and Westhof, 2015), and the prediction tasks were usually performed separately. Thus, the small sizes of individual binding data sets prevent the applications of deep learning techniques. In fact, most biological molecules are organic molecules, and the similarities in physic-chemical properties enable the sharing of interaction patterns. For example, the combined inputs of DNA-and RNA-binding residues into the same learning system have been proven to improve the predictions through the support vector machine techniques (Zhang X, 2016) (Su, et al., 2019) and the artificial neural network (Zhang, et al., 2012). However, these studies have ignored the differences between DNA and RNA molecules, and no study has yet been performed to combine binding information with other molecular types. For this purpose, the multi-task learning provides a promising framework that learns shared information through common networks while retaining the task-related output layers (Rich Caruana, 1997). The shared networks between multiple similar tasks enable a bigger training set that could maintain a larger network.

In this study, we designed a multi-task network architecture (namely MTDsite) to simultaneously predict respective binding residues with DNA, RNA, carbohydrate, and peptide molecules. The shared networks among all tasks can help learn common representations and thereby obtain relatively strong abstracting capabilities, and we used the LSTM as our Shared network to collect the information of long-range residues in the protein chain. At the same time, four small specific sub-networks were respectively trained for four individual types to extract individual properties. The benchmark tests indicated the employing of multi-task learning leads to averagely 3.6% improvements over state-of-the-art methods when measured by the area under the receiver operating characteristic curve (AUC).

## 2. Methods

### 2.1 Benchmark Datasets

We evaluated our method by using the previously curated training and testing datasets. The datasets include the protein binding with DNA, RNA, peptide, and carbohydrate, where a residue was defined as a binding residue if it contains at least one atom within 3.5Å from its binding partner. The sequence identities between the proteins in the training and test sets are less than 30% according to BLASTCLUST (Johnson, 2008). Table 1 lists the datasets, with details as:

**Table 1.** Summary of the benchmark datasets

| *Types* | *Traning* |  | *Test* |  | *References* |
| --- | --- | --- | --- | --- | --- |
|  | Chains | BD/ nonBD | Chains | BD/ nonBD |  |
| *DNA* | 309 | 6832/58270 | 47 | 875/8231 | (Su, et al., 2019) |
| *RNA* | 157 | 4627/30052 | 17 | 409/5448 | (Su, et al., 2019) |
| *Peptide* | 1115 | 14953/251708 | 125 | 1716/29154 | (Zhou, 2016) |
| *CBH* | 100 | 1028/25958 | 49 | 508/13230 | (Ghazaleh Taherzadeh, 2016) |

#### The DNA and RNA datasets

The datasets were collected from a recent study (Yan and Kurgan, 2017), where the training and testing datasets are 309 and 47 chains for DNA, and 157 and 17 chains for RNA, respectively.

#### The peptide dataset

The dataset was downloaded from a recent study (Taherzadeh G, et al., 2016), where protein–peptide complex structures were extracted from the BioLip protein–ligands database (Yang, et al., 2013) with peptides as the ligands derived from the Protein Data Bank (PDB). The dataset includes 1115 proteins as the training set, and 125 proteins as the independent test set that were randomly split by the previous study.

#### The carbohydrate dataset (CBH)

The dataset was downloaded from a recent study (Zhao, 2014) that includes 157 chains for training and 17 chains for test. The dataset was originally derived from the PROCARB (Malik, et al., 2010) by keeping only high resolution (<3Å) complex structures.

### 2.2 Input Features

Our input includes totally 54 features that are composed of G-PSSM (20 features), G-HHM (20), and G_SPD3 (14), as detailed below:

#### G-PSSM

Evolutionarily conserved residues are considered to be the same or similar residues maintained between species by natural selection. They have important functional roles like acting as binding sites. In this study, we employed the position specific scoring matrix (PSSM) which is a 20*L dimensional matrix (where L is protein length) generated from PSI-BLAST with E-value threshold of 0.001 in three iterations.

#### G-HHM

The hidden Markov models (HMMs) have been successfully used in protein structure prediction (Remmert, 2012) that assumed a Markov process with unobserved states, and the profile HMMs accomplish the protein structure prediction task well based on the HMM-HMM alignment. The sequence alignment generated by HHblits has been found with higher accuracy than by PSI-BLAST (Stephen F. Altschul, 1997). Here, HMM profile was generated by HHblits that compared to the query sequence with the proteins in the uniprot20_2015_06 database

#### G-SPD3

Predicted structural properties by SPIDER3

The structural properties were predicted by SPIDER 3.0 (Rhys Heffernan, 2018), including:

#### ASA (2)

The accessible surface area (ASA) means the surface area of a biomolecule accessible to a solvent, which reflects the functional importance of residues. Here, ASA of each residue was obtained by SPIDER 3.0, and the ASA was predicted as relative ASA named as rASA. We also computed the average rASA of the residue and its four adjacent residues.

#### Torsional angles (8)

The backbone torsional angles are composed of Φ, ψ, and ω that are used to describe local backbone structure of a protein. The ω was not used here because it is usually at 180° due to the planarity of the pep-tide bond. The angles between neighboring residues include θ and τ. Here, we let θ be the angle between Cαi-1-Cαi-Cαi+1, and τ be the dihedral angle rotated about the Cαi-Cαi+1 vector to compute the feature. We extracted the features by using cosine and sine values of the four angles, totally 8 features.

#### CN (1)

CN is the number of residues within a distance cutoff to a given residue in three-dimensional space.

#### HSE (3)

Half-Sphere Exposure (HSE) is an extension of CN, which considers directions of two residues in a top and bottom half of the sphere. Two methods were used to define the plane separating the upper and lower hemisphere, which included HSEα based on the neighboring Cα–Cαdirectional vector and HSE β based on the Cα–Cβ directional vector. In this study, we used the HSE α to describe the feature. For CN and HSE, residue distance is defined as the distance between Cα atoms with a 13 Å cutoff.

### 2.3 Performance evaluation

The performance of methods was measured by the number of correct classified and the number of misclassified instances using the terms below:

**TP:** number of actual binding residues predicted as binding sites
**TN**: number of actual non-binding residues predicted as non-binding sites
**FP:** number of actual non-binding residues incorrectly predicted as binding sites
**FN**: number of actual binding residues incorrectly predicted as non-binding sites

We evaluated the performance of our proposed prediction method in terms of the Matthews’ Correlation Coefficient (MCC), Accuracy, Receiver Operating Characteristic (ROC) curve as:

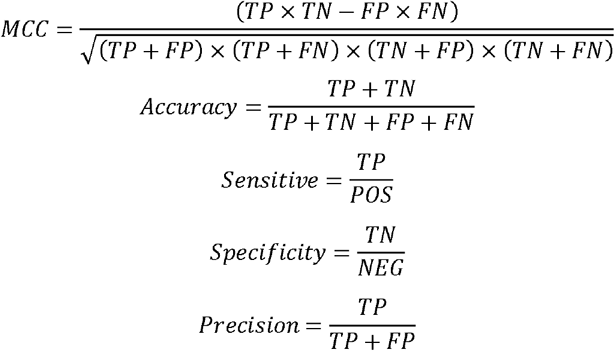
where POS is the number of known binding residues, and NEG is the number of known non-binding residues. MCC varies between 0 and 1, with 1 representing correct prediction for all residues, and 0 by random. Additionally, the Area Under the Curve (AUC) was adopted as our primary evaluation index because of our unbalanced datasets, i.e., the much less numbers of positive than negative samples.

### 2.4 MTDsite architecture

As shown in Fig 1, the deep learning network in this method consists of two parts. The first part is a shared Bi-directional Long Short-term Memory (BiLSTM) network called shared network (referred to shared BiLSTM), which was used to extract common information from different binding molecules. The second part is four individual networks composed of full connection layer. The predictive fraction of protein-molecules binding residues can be obtained from this part.

**Fig 1.**
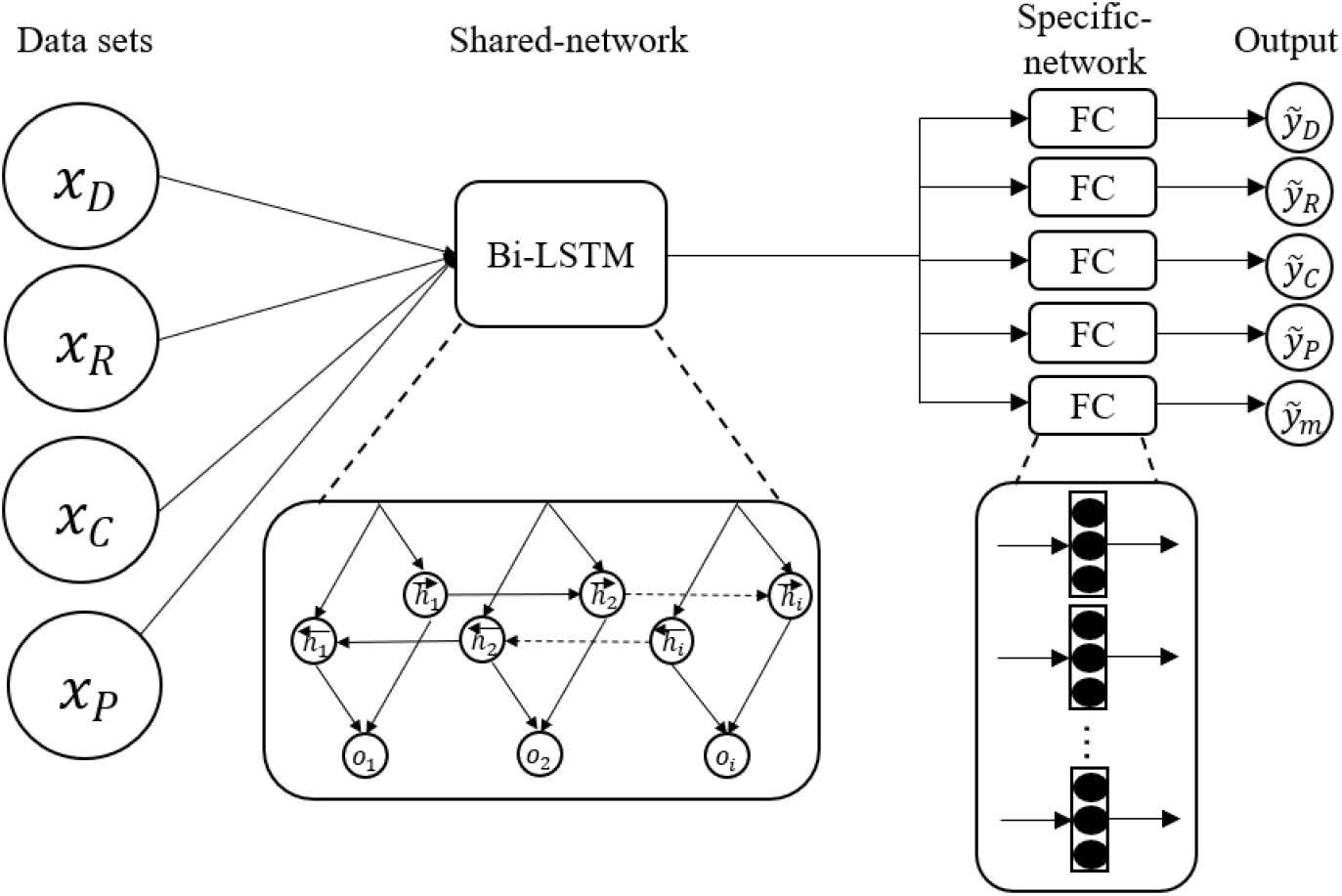
The general architecture of the MTDsite

#### Bi-LSTM network

Long short-term memory (LSTM) is a recurrent neural network (RNN) architecture (Hochreiter and Schmidhuber, 1997). The LSTM network was shown to have better performances than traditional RNN in processing, classifying, and predicting time series when there are indefinitely long separations between important events (Zhiheng Huang, 2015). This is the main reason why LSTM outperforms alternative Hidden Markov Models and other sequence learning methods in numerous applications (Yequan Wang, 2016).

In the prediction of binding residues, extracting information between long range residue pairs is important for constructing an accurate model. Traditional machine learning methods, such as Xgboost and SVM, extract feature information of adjacent residues by creating slide-window. In comparison, LSTM collects information on adjacent residues by establishing various ‘gates’, which saves time on training and adjusting parameters.

BiLSTM is a combination network of BRNN and LSTM by stacking multiple BiLSTM-RNN hidden layers together to build a deep bidirectional LSTM-RNN. The output of one layer is used as the input of the next layer. The hidden state sequence, *h^n^*, consist of forward and backward sequence 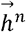 and 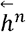
, iteratively computed from *n* = 1 to N and t = 1 to T as follows:

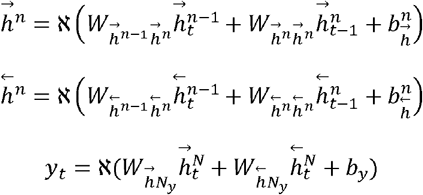

where y = (*y*_1_, *y*_2_, *y*_3_, …, *y_t_*, …, *y_T_*) is the output, ℵ is the activation function

#### Multi-task learning

A neural network is a non-linear classifier that performs repeated linear and non-linear transformations on the input. Let xi represent the input of i-th layer of the network (where x0 is just the feature vector). The transformation performed as:

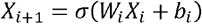

where *W_i_* and *b_i_* are the weight matrix and bias value for the *i*-th layer respectively, and *σ* is a nonlinear function. After N conversions, the final layer of the network *X^N^* is then fed to a simple linear classifier (the softmax in our method) to predict the probability of input *x*_0_ in label j:

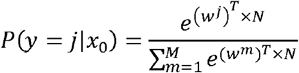

where M is the number of possible labels (here M = 4) and *w*^1^, *w*^2^, …, *w_M_* are weight vectors. The above parameters *W_i_*, *b_i_* and *w^m^* were parameters learned during training by the back-propagation algorithm. A multitask network attaches N SoftMax classifiers, one for each task, to the final layer *x_L_*. Here, we defined a “task” as classifier associated with a particular task, and one particular binding data set in our collection.

#### MTDsite networks

The networks consist of two parts, the shared network and four specific networks for individual tasks. The input of shared network is a 54 × L (L is the length of a protein and 54 is the number of the features) feature matrix. Through two shared LSTM hidden layers, a 2×L scoring matrix can be obtained, which is used as input for the next part.

As different tasks have specific properties, the second part is four independent fully-connected networks specifically trained for four different tasks. The four specific networks have identical structures (layer and neuron sizes). For each task, only the corresponding specific network will be updated, with the left three specific networks unchanged.

### 2.5 Cross-Validation and Independent Test

The cross validations and independent tests were employed to evaluate the robustness and performance of the method. The training data set was evenly divided into ten pieces (folds) at random. In each round, nine folds were employed for training and the remaining fold was used for test. This process was repeated for 10 times so that each fold has been tested once, and all outputs were collected to compute the performances. Based on the optimal hyper-parameters, a model trained by using the whole training set was then tested on the independent test set.

### 2.6 Model selection and Parameters optimization

During the optimization of the MTDsite models and hyperparameters, we only randomly selected 1/10 of the training samples, and selected the optimal parameters with the highest AUC value. We didn’t use the 10-fold cross validation to select as it will take 10 times of training costs. Finally, we used the ELU and Cross-Entropy as the network activation and loss functions in order to improve convergence speed and accuracy. The final optimal parameters of EPOCH and LR were 21 and 0.001, respectively, the number of hidden layers is 2, and the hidden nodes for shared network were both set as 128. The specific networks were all set as simple, single-layer, fully connected networks with 64 hidden nodes. These hyper-parameters (LR, the number of hidden layers and the number of hidden nodes) were determined by the GridSearchCV () function in python sklearn library. Due to the different lengths of all protein sequences, batch-size was set to 1 to avoid the decrease in accuracy caused by padding, though it reduces the training speed of the network. After the optimal parameters were decided, the models were evaluated by the cross-validations on the training sets, and independent tests on the test sets.

### 2.7 MTDsite-single Models

For a direct comparison, we trained MTDsite-single models like traditional ways by independently inputting single binding type of training data, and thus four models were obtained for four binding types, respectively. If not specifically mentioned, the model trained on one binding type will be employed to test the corresponding type. The architecture of MTDsite-single is a two-layer BiLSTM network, which is the same as the shared network of MTDsite. Similarly, the hyperparameters were also optimized on the training sets. The final EPOCH parameters are 17, 19, 26, and 28 for DNA, RNA, peptide and carbohydrate-binding models, and the learning rate is 0.001.

## 3. Results

### 3.1 Performances of MTDsite on the 10-fold cross validations and independent tests

Table 2 shows the performances of MTDsite measured by AUC, MCC, sensitivity, and specificity in prediction of DNA-, RNA-, peptide-, and carbohydrate-binding residues. By the 10-fold cross validation, MTDsite achieved AUC values of 0.866, 0.857, 0.760, and 0.779 for the four binding types, respectively. The predictions of DNA-and RNA-binding residues have relatively greater AUC values, likely because DNA and RNA are negatively charged and thus the binding residues are relatively easier to predict. The independent tests have obtained essentially the same AUC values with differences of only 0.003~0.024 for the four tests, indicating the robustness of our models. Fig 2 shows the ROC curves by the cross validations and independent tests on four binding types, respectively. The performances were also confirmed by the consistent maximum MCC values between the 10-fold cross validations and independent tests, with differences of 0.003~0.021. At the thresholds with the maximum MCC values, our models have high specificities while relatively low sensitivities due to the much greater numbers of non-binding residues (negatives) than the binding residues (positives).

**Figure 2.**
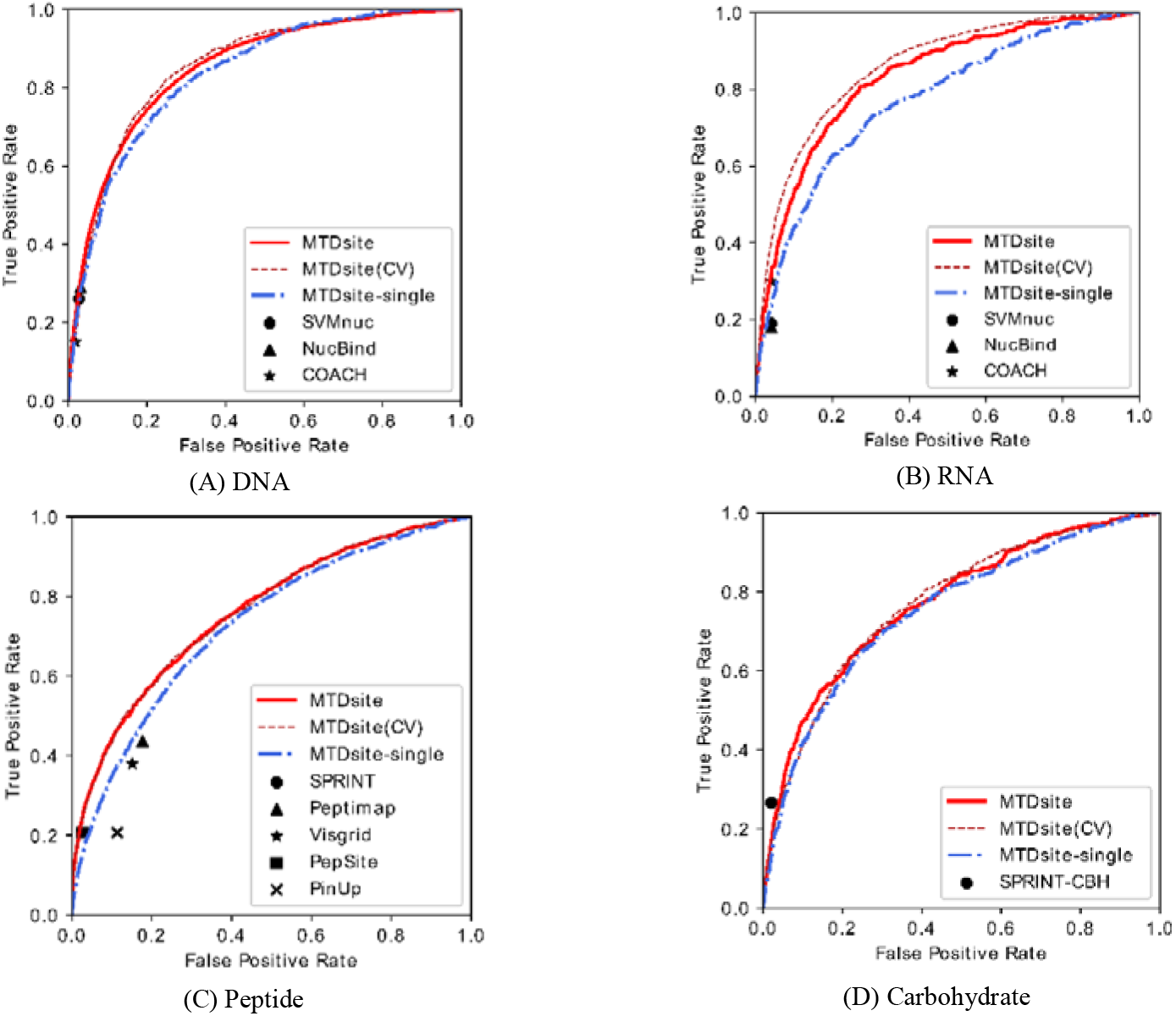
The receiver of characteristic curves for MTDsite (10 fold cross-validations and independent tests) and MTDsite-single on the independent tests for (A) DNA, (B) RNA, (C) peptide, and (D) carbohydrate datasets. The reported results by other methods were labelled on respective plots.

**Table 2.** Method performance on 10-fold cross validation and independent test

| **Dataset** |  | **AUC** | **MCC** | **Sen** | **Spe** |
| --- | --- | --- | --- | --- | --- |
| **DNA** | CV | 0.866 | 0.401 | 0.691 | 0.877 |
|  | Test | 0.852 | 0.397 | 0.684 | 0.873 |
| **RNA** | CV | 0.857 | 0.369 | 0.635 | 0.936 |
|  | Test | 0.836 | 0.361 | 0.612 | 0.932 |
| **Peptide** | CV | 0.760 | 0.304 | 0.346 | 0.956 |
|  | Test | 0.758 | 0.299 | 0.344 | 0.953 |
| **CBH** | CV | 0.779 | 0.276 | 0.428 | 0.957 |
|  | Test | 0.776 | 0.255 | 0.414 | 0.954 |

We evaluated the contributions of individual feature group by using only single feature group or excluding one feature group from all features. As shown in Table 3, when individual feature group was used in the prediction, G-PSSM, the evolution features produced by PSIBLAST, yielded the greatest values in regard with the average values of both AUC and MCC. G-HHM, another feature group produced by HHblits, yielded slightly lower AUC and MCC values. G-SPD3, the structural feature group produced by SPIDER3 package, yielded significantly lower AUC values in average. These results suggest the importance of evolution information for protein binding, consistent with previous findings (Hong Su, et al, 2018). When excluding individual feature group, the removal of G-PSSM caused the largest decreases in the average values of both AUC and MCC, again indicating its most important role. Though the removal of the G-SPD3 feature group caused the smallest decrease, the difference is significant (P=0.001) according to the paired t-test. The decreases are small likely because the G_SPD3 features were derived from the PSSM and HHM profiles, and our neural networks could partly catch the structural information from the two profiles.

**Table 3.** The average AUC and MCC values on four independent datasets by employing or excluding individual feature group from the MTDsite model.

| **Feature Groups<sup>a</sup>** | **AUC** | **MCC** | **Feature Groups<sup>b</sup>** | **AUC** | **MCC** |
| --- | --- | --- | --- | --- | --- |
| - | - | - | MTDsite | 0.806 | 0.328 |
| G-PSSM | 0.742 | 0.247 | -G-PSSM | 0.757 | 0.281 |
| G-HHM | 0.739 | 0.244 | -G-HHM | 0.767 | 0.302 |
| G-SPD3 | 0.61 | 0.201 | -G-SPD3 | 0.794 | 0.320 |

<sup>a</sup> Performances based on individual feature groups;
<sup>b</sup> Performances by excluding individual feature groups.

### 3.2 Contributions by the shared networks

To evaluate the contributions of the networks shared by different binding types, we trained four MTDsite-single models for comparison, each with single type of binding data. As shown in Fig 2, the ROC curves of MTDsite-single are consistently below those of MTDsite by independent tests on four binding types. As a result, the AUC values by MTDsite-single are 3.6%, 3.2%, 3.3%, and 4.0% lower than those by MTDsite for DNA, RNA, peptide, and carbohydrate, respectively (Table 4). The difference of ROC curves between MTDsite and MTDsite-single reflect the improvement affected by the data set fusion and multi-task learning. When measured by MCC, the MTDsite-single are 4.7%, 36.7%, 37.2%, and 15.4% worse than the MTDsite.

**Table 4.**
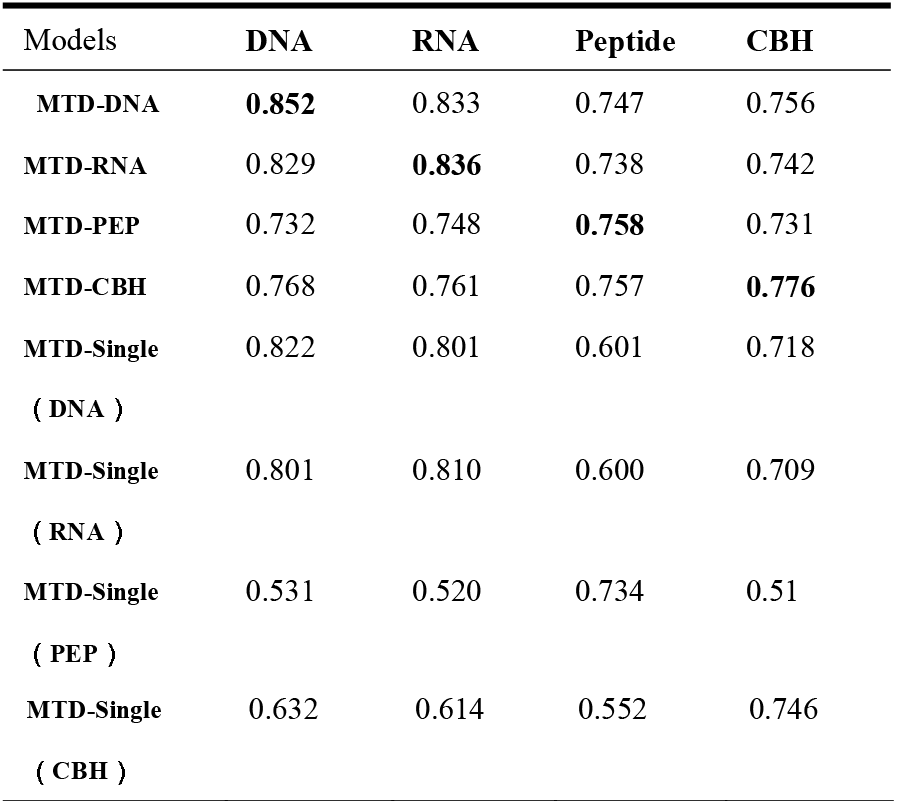
Comparison of AUC values of different MTDsite models for different binding molecules on Independent test

| Models | **DNA** | **RNA** | **Peptide** | **CBH** |
| --- | --- | --- | --- | --- |
| **MTD-DNA** | **0.852** | 0.833 | 0.747 | 0.756 |
| **MTD-RNA** | 0.829 | **0.836** | 0.738 | 0.742 |
| **MTD-PEP** | 0.732 | 0.748 | **0.758** | 0.731 |
| **MTD-CBH** | 0.768 | 0.761 | 0.757 | **0.776** |
| **MTD-Single** | 0.822 | 0.801 | 0.601 | 0.718 |
| ( DNA ) |  |  |  |  |
| **MTD-Single** | 0.801 | 0.810 | 0.600 | 0.709 |
| ( RNA ) |  |  |  |  |
| **MTD-Single** | 0.531 | 0.520 | 0.734 | 0.51 |
| ( PEP ) |  |  |  |  |
| MTD-Single | 0.632 | 0.614 | 0.552 | 0.746 |
| ( CBH ) |  |  |  |  |

We further performed cross tests by using the classifier trained from one binding type to test other types. As shown in Table 4, on prediction of DNA and RNA binding residues, similar performances have been achieved by the MTD-single (DNA) and MTD-single (RNA), indicating a similarity between DNA-and RNA-binding sites. This also explains why predictions could be improved by a simple combination of DNA and RNA binding residues in the previous studies. By comparison, there are significant decreases on other cross tests, among which the MTD-single (PEP) made almost random predictions on other three binding types although the peptide-binding dataset used for training has the biggest sample size. Interestingly, the MTD-single (DNA) and MTD-single (RNA) models produced good predictions on the carbohydrate-binding dataset: only 3.9% and 5.2% lower AUC than MTD-single (CBH). This is likely because carbohydrate has common properties with the desoxyribose and ribose respectively contained in DNA and RNA molecules. By comparison in the MTDsite, although the specific networks have always produced the best predictions for their respective binding types, other specific networks in the MTDsite could also achieve reasonable predictions. For example, on the prediction of peptide-binding residues, MTDsite (CBH) achieved essentially the same AUC value as the MTDsite (PEP), indicating its potential generality to other binding types.

Fig 3 shows a direct comparison of MTDsite and MTDsite-single. Among the 237 protein chains from four independent datasets, MTDsite significantly outperforms MTDsite-single with P-value of 0.003 according to the paired t-test, where MTDsite have greater AUC value for 162 (68%) chains.

**Figure 3.**
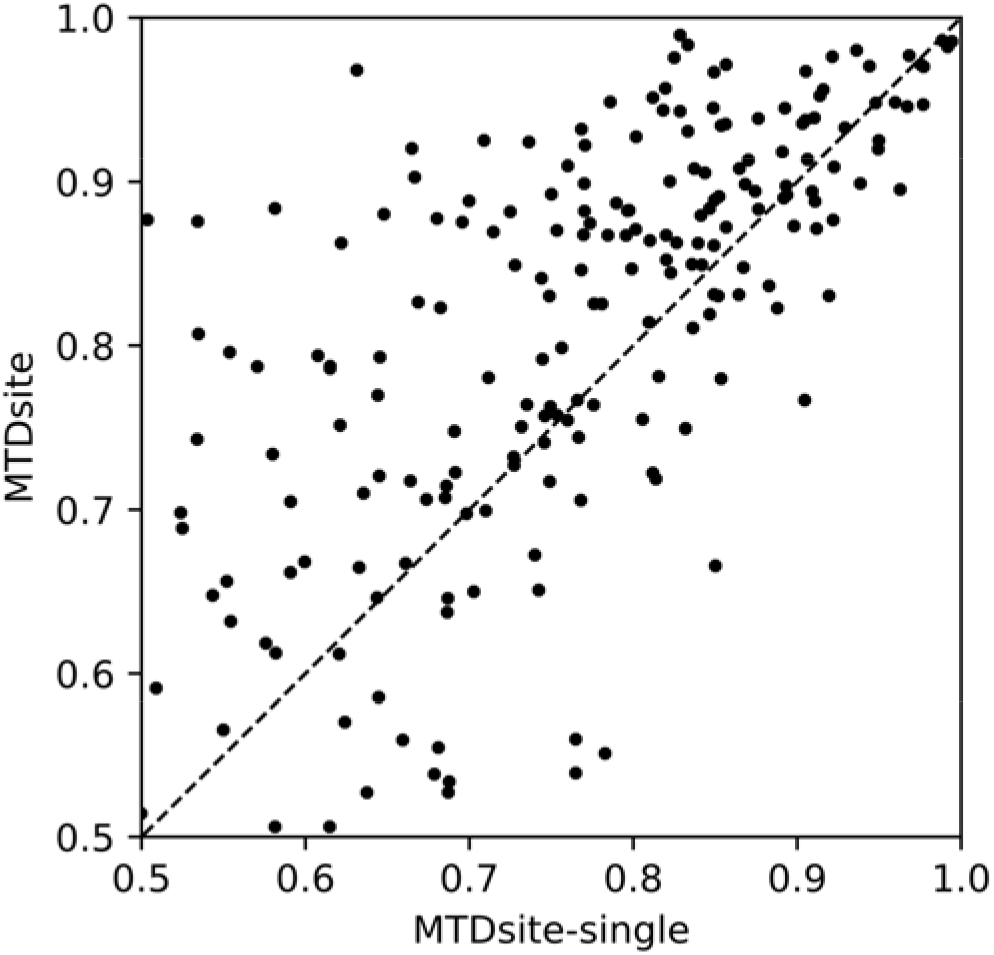
The comparison of AUC values by MTDsite and MTDsite-single on independent dataset, each point representing one protein chain.

### 3.3 Comparisons with other methods

MTDsite was compared with other methods on the independent test sets respectively for four binding types. As shown in Table 5, MTDsite achieved the highest AUC values through all four types of independent datasets, which are 2.2%, 4.4%, 6.6%, and 0.52% higher than other best methods for DNA, RNA, peptide, and carbohydrate datasets, respectively. The improvements are mostly contributed by the multi-task learning as the MTDsite-single models without using multi-task learning yielded close and mostly lower performances than other state-of-the-art methods. Fig 2 also indicated that all other methods are mostly below the ROC curves by MTDsite in all four tests. The only exception is the SPRINT-CBH method, previously developed by our group for the carbohydrate-binding prediction, which is marginally above the ROC by MTDsite, though MTDsite achieved a higher AUC value. Here, the lower MCC by MTDsite is likely because MTDsite has been optimized for the greatest AUC values while SPRINT-CBH was optimized for MCC values. Additionally, the carbohydrate dataset shows a much larger ratio between the number of negatives and the positives in the test set (26.0 on the carbohydrate compared to 8.5, 6.5, and 16.8 on the DNA, RNA, peptide datasets, respectively), and thus the shared network wasn’t well optimized for the carbohydrate dataset.

**Table 5.** Comparison with other methods on the independent test sets based on AUC, ACC, MCC.

Sensitivity and Precision
|  | Method | AUC | MCC | ACC | Sen | Pre |
| --- | --- | --- | --- | --- | --- | --- |
| **DNA** | BindN+ | 0.806 | 0.256 | NA | NA | NA |
|  | SVMnuc | 0.833 | 0.336 | NA | 0.26 | 0.51 |
|  | NucBind | 0.834 | 0.355 | NA | 0.29 | 0.50 |
|  | DRNApred | 0.757 | 0.232 | NA | 0.32 | 0.25 |
|  | COACH-D | 0.810 | 0.266 | NA | NA | 0.70 |
|  | MTDsite-single | 0.822 | 0.379 | 0.901 | 0.562 | 0.628 |
|  | MTDsite | **0.852** | **0.397** | **0.916** | **0.684** | **0.734** |
| **RNA** | BindN+ | 0.738 | 0.219 | NA | NA | NA |
|  | SVMnuc | 0.784 | 0.205 | NA | 0.19 | 0.26 |
|  | NucBind | 0.801 | 0.210 | NA | 0.18 | 0.29 |
|  | DRNApred | 0.687 | 0.056 | NA | 0.09 | 0.08 |
|  | COACH-D | 0.711 | 0.356 | NA | 0.30 | **0.45** |
|  | MTDsite-single | 0.810 | 0.264 | 0.945 | 0.455 | 0.345 |
|  | MTDsite | **0.836** | **0.361** | **0.931** | **0.612** | 0.351 |
| **Peptide** | VisGrid | NA | 0.224 | 0.609 | 0.38 | NA |
|  | PinUp | NA | 0.0071 | 0.547 | 0.207 | NA |
|  | PepSite | NA | 0.141 | 0.598 | 0.221 | NA |
|  | Peptimap | NA | 0.262 | 0.629 | 0.436 | NA |
|  | SPRINT | 0.711 | 0.279 | 0.662 | **0.642** | NA |
|  | MTDsite-single | 0.734 | 0.218 | 0.947 | 0.32 | 0.29 |
|  | MTDsite | **0.758** | **0.299** | **0.956** | 0.344 | **0.366** |
| **CBH** | SPRINT-CBH | 0.772 | **0.285** | 0.961 | **0.223** | NA |
|  | MTDsite-single | 0.746 | 0.221 | 0.958 | 0.374 | 0.177 |
|  | MTDsite | **0.776** | 0.255 | **0.963** | 0.209 | 0.171 |

As a result, the MCC values of MTDsite are 11.8%, 1.4%, and 7.2% percent higher than other methods for DNA, RNA, and peptide, respectively. At the same time, MTDsite achieved the highest accuracy, sensitivity, and precision in most cases. It should also be noticed that PepSite and Peptimap are structure-based methods. Though MTDsite used sequence-based information only, our method still achieved better performances.

### 3.4 Case study

As an example, we demonstrated the prediction on a restriction-modification controller DNA-binding protein (PDBid: 3s8qB). As shown in Fig 4, MTDsite predicted 15 binding residues that include 14 residues are truly binding. In comparison, MTDsite-single predicted 14 binding residues including 13 truly binding residues. Thus, total accuracies of two methods are 97% and 94%, respectively. The difference of MTD-single in prediction performance on different data sets showed that our method could distinguish the binding sites of different molecules on proteins, and also proved that cross-prediction had less impact on MTDsites.

**Figure 4.**
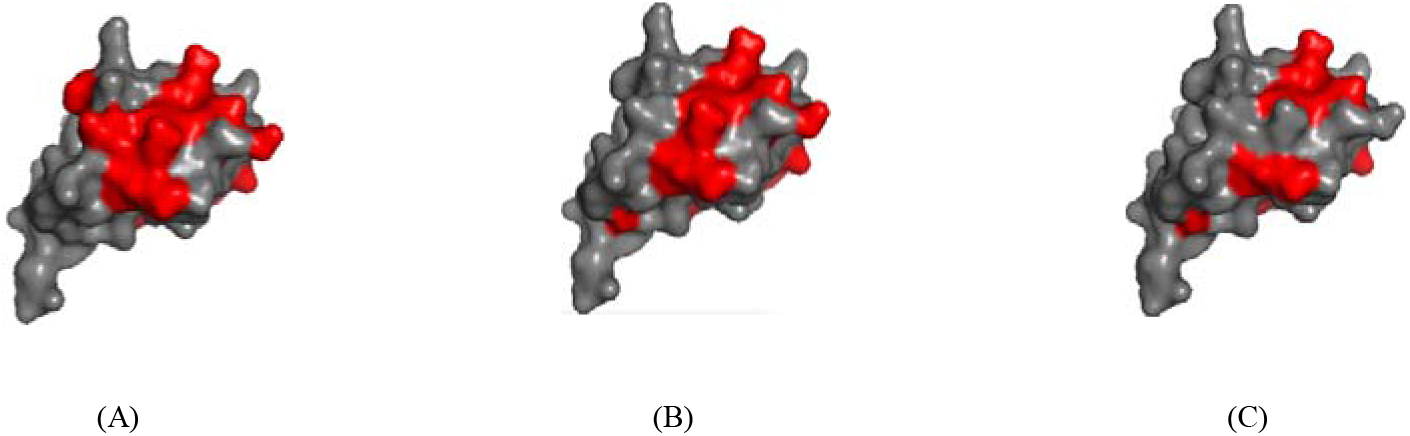
(A) The actual binding residues, and the predicted binding residues by (B) MTDsite and (C) MTDsite-single for a Restriction-Modification controller DNA-binding protein (PDBid: 3s8qB).

## 4. Conclusion and Discussion

We have developed a new architecture to use multiple-task learning for predicting binding residues. To our best knowledge, this is the first attempt of the strategy for prediction of binding residues. The sharing between DNA, RNA, peptide, and carbohydrate-binding information was indicated to improve the predictions of binding residues for all four types. Our method was indicated robust by the consistent performances between the cross validations and independent tests. The method was proven to outperform state-of-the-state methods. Such strategy was expected to extend to other tasks like predictions of ligand-and lipid-binding residues. The framework is also promising for other similar tasks, such as protein function prediction, modification sites of protein or DNA/RNA, and predicting properties for chemical compounds.

As this study focused on predicting binding residues from sequences, the prediction from protein structures (from experiments or structural modeling) was expected to further advance the predictions. This has been proved by our previous studies (Taherzadeh, et al., 2017).

The current sharing of information was simply based on a port of shared network. The performances were hurt by the differences between the binding types, for example, different ratios of positives and negatives, the preference of positive-charge residues in the DNA/RNA binding. Recently, there is a significant advance in the transferred learning techniques (José Juan Almagro Armenteros, 2019). We will employ these recent techniques in future studies.

## 5. Acknowledgement

This study was supported in part by the National Key R&D Program of China (2018YFC0910500), National Natural Science Foundation of China (U1611261, 61772566, and 81801132), Guangdong Frontier & Key Tech Innovation Program (2018B010109006, 2019B020228001) and Introducing Innovative and Entrepreneurial Teams (2016ZT06D211).

## Notes

http://biomed.nscc-gz.cn/server/MTDsite/

